# Retinal Electrophysiological Patterns in Alzheimer’s Disease: A Multi-Domain Signal Processing Framework for Non-Invasive Biomarker Discovery Using a Portable ERG Device

**DOI:** 10.64898/2026.05.20.726572

**Authors:** Joel A. Barría, Andrea Slachevsky, Adrián G. Palacios, Leonel E. Medina

## Abstract

Alzheimer’s disease (AD) is a neurodegenerative disorder affecting more than 55 million people worldwide, with a diagnosis that remains predominantly clinical and frequently delayed. The electroretinogram (ERG) offers a non-invasive electrophysiological method for detecting retinal dysfunction associated with neurodegeneration; however, it remains unclear whether robust and reliable candidate biomarkers can be extracted from ERG signals beyond conventional amplitude- and latency-based parameters. Here we present a pilot study of a multi-domain signal processing framework applied to ERGs recorded from 46 participants (20 AD patients, 26 controls) with a handheld device (RETeval, LKC Technologies) using sinusoidal (1–50 Hz) and photopic ISCEV protocols. Five complementary techniques were implemented: (i) multiscale fuzzy entropy (MSFuzzyEn); (ii) FFT harmonic analysis; (iii) stimulus-response wavelet time-frequency coherence (WTC); (iv) a novel inter-cycle lag variant of sample entropy (SampEn_*T*_), introduced to isolate cycle-to-cycle retinal response consistency independently of stimulus periodicity; and (v) discrete wavelet transform (DWT) for energetic extraction of oscillatory potentials (OPs). Univariate comparisons (Mann-Whitney, Cliff’s *δ*, Benjamini-Hochberg FDR) identified seven significant candidate biomarkers (*q <* 0.05), five with large effect size: AUC_fast_ (|*δ*| = 0.546, *q* = 0.009), Slope_very-slow_ (|*δ*| = 0.554, *q* = 0.007), *R*_14*f*_ (|*δ*| = 0.515, *q* = 0.031), SampEn_*T*_ (|*δ*| = 0.504, *q* = 0.019) and WTC_*R*,mean_ (|*δ*| = 0.531, *q* = 0.023); and two with medium effect size (OP_amp_sum, band_snr). A logistic regression classifier combining three candidate biomarkers, validated by leave-one-out cross-validation, achieved ROC-AUC = 0.858, sensitivity = 70.0% and specificity = 88.5% (*n* = 46). These proof-of-concept results demonstrate that multi-domain ERG analysis captures retinal temporal dysfunction signatures in AD that are inaccessible to standard clinical analysis, supporting further investigation of portable ERG devices as a source of non-invasive candidate biomarkers for early AD detection.

## 1. Introduction

Alzheimer’s disease (AD) is the most common cause of dementia, affecting more than 55 million people worldwide, a figure projected to triple by 2050 [1]. Current diagnosis relies on cerebrospinal fluid (CSF) biomarkers and positron emission tomography (PET), procedures that are invasive, expensive, and inaccessible in most clinical settings [2, 3]. The need for non-invasive, affordable, and scalable biomarkers capable of detecting AD at early stages is a recognized global public health priority.

### 1.1. The retina as a candidate tissue for biomarkers

The retina shares embryological and structural properties with the brain: both tissues originate from the neuroectoderm, share the blood-brain barrier, and exhibit homologous vascularization patterns and neuronal circuits [4]. This anatomical and functional continuity makes it a particularly relevant tissue for the study of neurodegeneration. Both amyloid-*β* and tau deposits have been detected in retinal tissue of AD patients, and histological changes—including loss of retinal ganglion cells (RGC) and reduction of nerve fiber layer thickness—may precede manifested cognitive symptoms [2, 5, 6].

The electroretinogram (ERG) provides an objective and quantitative measure of retinal electrical activity, accessible non-invasively with skin electrodes. Recent studies have shown that the ERG detects RGC dysfunction in preclinical AD through the photopic negative response (PhNR) [7], with ROC areas under the curve comparable to those of molecular biomarkers in at-risk populations. However, clinical ERG analysis is primarily based on the amplitude and implicit time of the main waves, namely, a, b, and PhNR, discarding the enormous wealth of information contained in the complete temporal structure of the signal.

### 1.2. The ERG as an information-rich non-stationary electrophysiological signal

The full-field ERG is a complex electrophysiological signal whose non-stationary nature—variable in amplitude, phase, and spectral content—demands analytical tools that go beyond conventional clinical parameters (a, b, and PhNR waves).

In full-field sinusoidal stimulation paradigms, inner retinal circuits—bipolar, amacrine, and ganglion cells—introduce nonlinearities that manifest as harmonic components at multiples of the stimulus frequency, visible in the Fourier spectrum of the signal [8, 9]. The relative amplitude of these harmonics reflects the functional state of inner retinal circuits and is sensitive to pathological conditions such as diabetic retinopathy [10, 11], where alterations in nonlinearities precede conventional amplitude changes. Note, however, that even during sustained sinusoidal stimulation the ERG signal exhibits cycle-to-cycle variability and slow amplitude modulations attributable to retinal adaptation and fluctuations in synaptic coupling [12], classifying it as a non-stationary electrophysiological signal in the strict sense.

In standard ISCEV (International Society for Clinical Electrophysiology of Vision) photopic flash paradigms, the ERG signal also contains oscillatory potentials (OPs) superimposed on the b-wave, generated by amacrine cell circuits of the inner retina in the 60–160 Hz band [9, 13]. The discrete wavelet transform (DWT) has been shown to be the reference method for extraction and energetic quantification of OPs, allowing selective separation of contributions from the sub-bands at 20, 40, 80, and 160 Hz corresponding to the ON/OFF pathways and to fast and slow OPs [14, 15]. This time-frequency decomposition capability surpasses classical bandpass filtering, which does not preserve the temporal localization of components [16, 13].

Multiscale entropy measures have demonstrated their ability to detect alterations in the complexity of electrophysiological signals both in animal models of AD, and in EEG and MEG recordings of patients [17, 18, 19]. Multiscale fuzzy entropy (MSFuzzyEn) constitutes the most robust variant for signals of moderate length [20], since the use of fuzzy membership functions reduces sensitivity to noise and abrupt amplitude changes inherent in non-stationary signals such as the ERG [21]. Its systematic application to the human ERG in a clinical context constitutes an original contribution of the present work.

Time-frequency wavelet coherence (WTC) offers a complementary perspective: rather than quantifying the intrinsic complexity of the signal, it measures the synchrony between the ERG response and the reference stimulus, capturing the consistency of retinal coupling over time [22, 23]. Unlike classical spectral coherence, WTC does not assume stationarity and is appropriate for signals with dynamic properties [24]. Wavelet analysis has previously been applied to the ERG for time-frequency domain feature extraction, with results superior to classical amplitude analysis in both glaucoma [16] and neurodevelopmental disorders [15].

In this work we introduce and evaluate a multi-domain signal analysis framework applied to full-field ERGs recorded with the portable RETeval device (Figure 1), as a proof-of-concept pilot study. Specifically, we integrate five complementary analytical domains — multiscale fuzzy entropy (MSFuzzyEn), FFT harmonic analysis, time-frequency wavelet coherence (WTC), inter-cycle sample entropy, and discrete wavelet transform (DWT) for oscillatory potential extraction — into a unified processing pipeline for sinusoidal and ISCEV ERG signals, and provide preliminary evidence that metrics derived from each domain significantly discriminate AD patients from healthy controls with large effect sizes. A minimum-dimensionality logistic regression classifier evaluated by leave-one-out cross-validation (LOO-CV) combines three candidate biomarkers from independent domains, achieving preliminary diagnostic performance that, if replicated in larger cohorts, may be comparable to that of invasive methods. Additionally, medium-effect metrics — including DWT energy of oscillatory potentials and narrowband SNR — are identified as potential contributors to the pathophysiological characterization of retinal dysfunction in AD.

**Figure 1:**
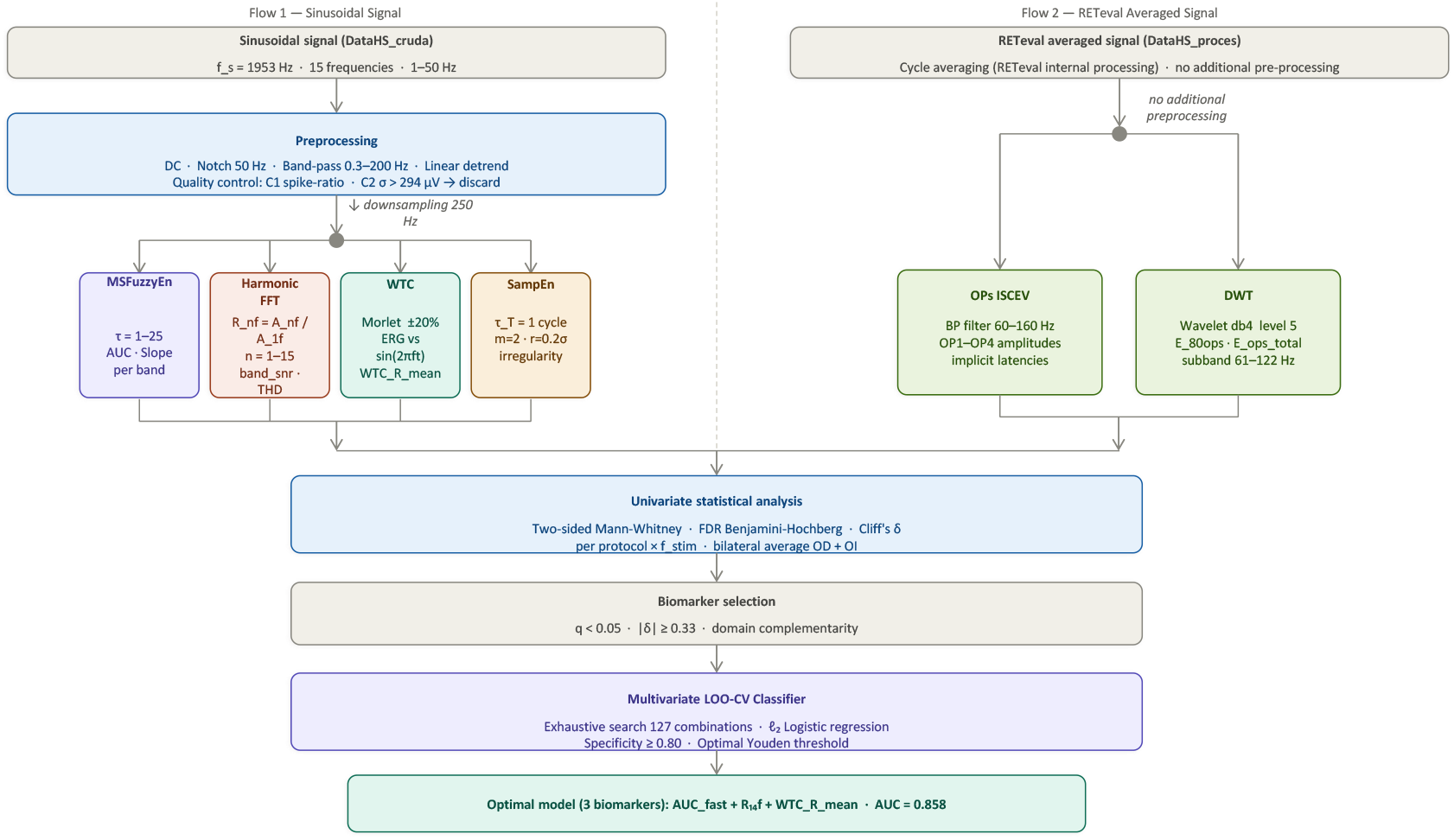
Multi-domain signal processing pipeline. Flowchart of the complete analysis chain, from ERG recording to LOO-CV classification. Raw signals (*f*_*s*_ = 1953.125 Hz) are split into two branches: **Branch I** (sinusoidal protocols) undergoes common preprocessing (DC removal, 50 Hz notch, 0.3–200 Hz bandpass, detrend) followed by extraction in four domains: MSFuzzyEn (*τ* = 1–25), FFT harmonic analysis, time-frequency wavelet coherence (WTC), and inter-cycle sample entropy (SampEn); **Branch II** (ISCEV protocol) directly uses the RETeval averaged signal for oscillatory potential extraction via DWT. Significant metrics from each domain converge into a logistic regression classifier evaluated by LOO-CV.

## 2. Materials and Methods

### 2.1. Participants

A total of 46 participants were included: 20 patients with Alzheimer’s disease dementia (AD; 9 M/11 F; age 69.6 ± 7.8 years, range 51–78) and 26 healthy controls (Ctrl; 2 M/24 F; age 66.3 ± 9.4 years, range 42–80). The two groups did not differ significantly in age (*p* = 0.28, Mann-Whitney *U* test). AD diagnosis followed NINCDS-ADRDA criteria and was confirmed by cognitive assessment (MMSE ≥ 15). Participants with significant systemic disease that could affect retinal function were excluded. One subject was excluded *a priori* due to acquisition artefacts (signal amplitude several orders of magnitude above physiological range), yielding *n* = 46. The sex distribution was unbalanced between groups (2 M/24 F in controls vs. 9 M/11 F in AD), which constitutes a limitation when generalizing our results.

All participants signed written informed consent. The study was approved by the Institutional Ethics Committee of Universidad de Santiago de Chile (Ethics Report No. 323/2023) and conducted in accordance with the Declaration of Helsinki. Measurements were carried out at the Memory Unit of Hospital del Salvador, Santiago, Chile.

### 2.2. ERG Recordings

Full-field ERGs were recorded bilaterally using the RETeval system (LKC Technologies Inc., Gaithersburg, MD, USA), a portable device that automatically compensates for pupil size to maintain constant retinal luminance, using skin electrode arrays (active, reference, and ground). Signals were digitized at *f*_*s*_ = 1953.125 Hz. Pharmacological dilation was not required. Three stimulation protocols were administered in fixed order to each eye:

1. **ISCEV photopic protocol:** three full-field photopic flash stimuli at the standard frequencies:

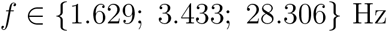

administered according to the ISCEV 2022 standard [25] on an adapted photopic background. The averaged signals generated by the RETeval device were used directly, without additional processing prior to oscillatory potential (OP) extraction.
2. **Sinusoidal 1–10 Hz protocol:** sinusoidal stimulation at 10 discrete frequencies:

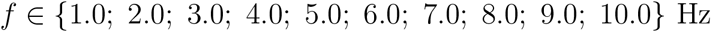

5 s per frequency at 100 cd/m^2^.
3. **Sinusoidal 10–50 Hz protocol:** sinusoidal stimulation at 5 discrete frequencies:

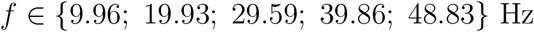

5 s per frequency at 100 cd/m^2^.

Together, the sinusoidal protocols cover the 1–50 Hz range, allowing characterization of the retinal temporal response from the low-frequency pathway (dominated by ON bipolar cells) to the high-frequency pathway (amacrine and ganglion cell circuits) [26]. The ISCEV protocol complements the analysis with the characterization of OPs under internationally standardized conditions.

### 2.3. Signal Preprocessing

ERG signals were processed in Python 3 with `NumPy` [27], `SciPy` [28], and `PyWavelets` [29]. Figure 1 summarizes both processing branches. Figure 2 illustrates representative ERG signals and biomarker profiles for subjects near the group median of each metric.

**Figure 2:**
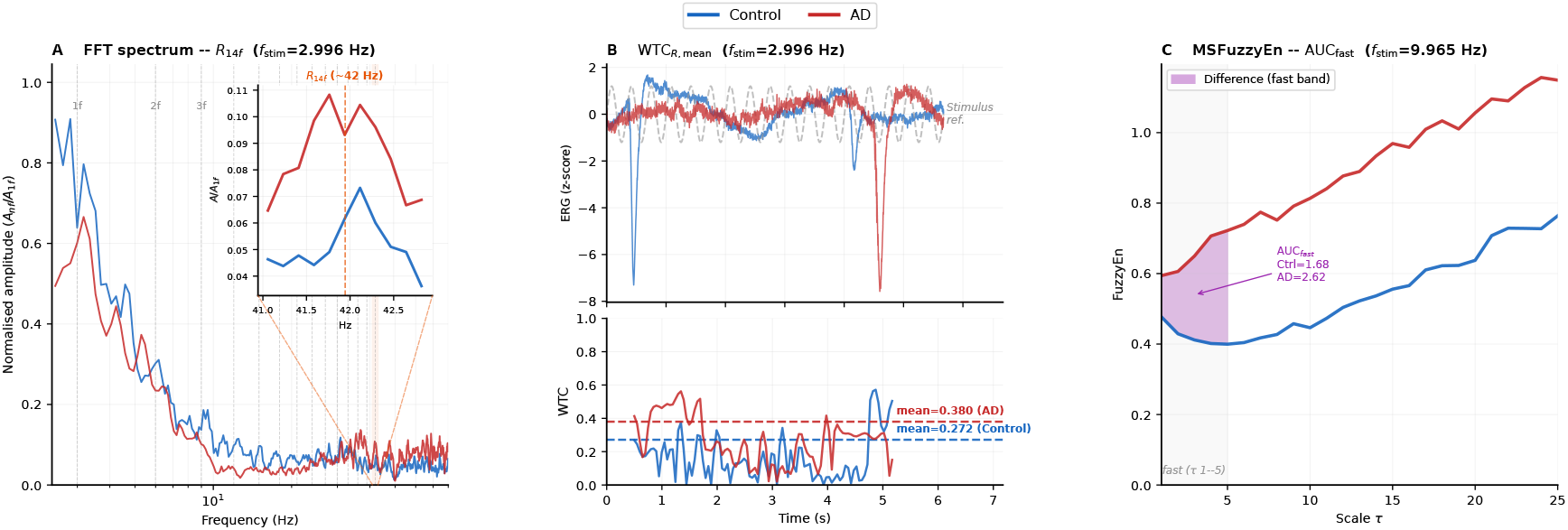
Representative ERG signals and candidate biomarkers of the parsimonious classifier (subjects near the group median). **(A)** Amplitude spectrum normalised by the fundamental frequency amplitude (*R*_*nf*_ = *A*_*nf*_ */A*_1*f*_), sinusoidal 1–10 Hz protocol, *f*_stim_ = 2.996 Hz (log scale). The inset magnifies the band around the 14th harmonic (*f*_14_ ≈ 41.9 Hz), where the AD subject presents a higher ratio *R*_14*f*_ than the control, consistent with the group medians (AD: 0.144; Ctrl: 0.098; |*δ*| = 0.515; *q* = 0.031). Spectral amplitudes are displayed with mild smoothing (moving average, window = 5 bins) for visualisation purposes only; all quantitative analyses were performed on the unsmoothed spectrum. **(B)** Preprocessed ERG signal (upper panel, z-score) and instantaneous time-frequency wavelet coherence WTC_*R*_(*t*) (lower panel) at *f*_stim_ = 2.996 Hz. Dashed lines indicate the temporal mean of WTC_*R*_ per group (AD: 0.388; Ctrl: 0.272; |*δ*| = 0.531; *q* = 0.023). **(C)** MSFuzzyEn profile as a function of temporal scale *τ* at *f*_stim_ = 9.965 Hz (10–50 Hz protocol). The shaded region (*τ* = 1–5) corresponds to the fast scales band over which AUC_fast_ is calculated (Ctrl = 1.68; AD = 2.62; |*δ*| = 0.546; *q* = 0.009). In all panels: blue = Controls (*n* = 26), red = AD (*n* = 20). *q*-values corrected by Benjamini-Hochberg FDR.

#### Branch I — Sinusoidal protocols (1–10 Hz and 10–50 Hz)

Raw signals recorded with RETeval were subjected to the following preprocessing pipeline, applied sequentially:

1. **DC component removal**.
2. **Notch filter** at 50 Hz (*Q* = 30, `filtfilt`).
3. **Bandpass filter:** 4th-order zero-phase Butterworth, [0.3; 200] Hz.
4. **Linear detrend** by least squares.

Preprocessed signals were used for feature extraction in four analytical domains: multiscale fuzzy entropy (MSFuzzyEn), FFT harmonic analysis, time-frequency wavelet coherence (WTC), and inter-cycle sample entropy (SampEn).

#### Branch II — ISCEV photopic protocol

Averaged signals internally generated by the RETeval device were used directly, without additional preprocessing. The discrete wavelet transform (DWT) was applied to these signals for extraction and energetic quantification of oscillatory potentials (OPs).

### 2.4. Quality Control and Artefact Rejection

Each preprocessed signal record was evaluated using two quantitative criteria. The first criterion, *C*_1_ = *p*_99_(|*x*_raw_|)*/*median(|*x*_raw_|) > 1.5, detects the presence of transient high-amplitude artefacts by comparing the 99th percentile of the absolute raw signal to its median: a ratio above 1.5 indicates that extreme values are disproportionately large relative to the central tendency of the signal [25]. The second criterion, *C*_2_ = *σ*(*x*_proc_) > 294 *µ*V, flags records with globally excessive noise by thresholding the standard deviation of the preprocessed signal; the threshold of 294 *µ*V corresponds to three times the 75th percentile (3 × *P*_75_) of the standard deviation distribution across all records in the dataset, a robust data-driven cutoff analogous to outlier detection methods based on the interquartile range [30]. Records failing either criterion were discarded prior to feature extraction. Of the 1,380 potential sinusoidal records, 16 were discarded in 6 subjects (4 Ctrl, 2 AD), all due to elevated *C*_2_, indicating globally noisy signals rather than isolated transient artifacts.

### 2.5. Multi-Domain Feature Extraction

#### 2.5.1. Domain I: Multiscale Fuzzy Entropy (MSFuzzyEn)

MSFuzzyEn quantifies the complexity of a signal at multiple temporal scales [21, 20]. Preprocessed signals were resampled to 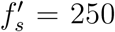 Hz via polyphase antialiasing decimation. For each scale *τ* ∈ {1, …, 25}:

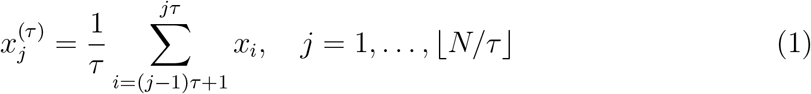

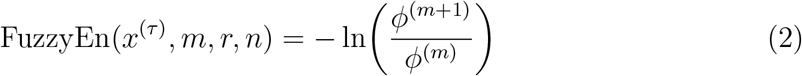

Parameters: *m* = 2, *r* = 0.15 *σ*(*x*), *n* = 2 [21].

The MSFuzzyEn profile was segmented into four scale bands following the coarse-graining framework [31]: fine scales *τ* ∈ [1, 5], medium-fine scales *τ* ∈ [5, 10], medium-coarse scales *τ* ∈ [10, 18], and coarse scales *τ* ∈ [18, 25] (Fig. 2C). From the MSFuzzyEn profile, the following features were extracted per scale band [31]: the area under the curve (AUC_band_), computed as the sum of FuzzyEn values over the band (Eq. 3); and the linear regression slope (Slope_band_), computed over the band (Eq. 4). Both features were computed independently for each of the four scale bands, yielding eight band-specific metrics: AUC_fast_, AUC_mid_, AUC_slow_, AUC_very-slow_, Slope_fast_, Slope_mid_, Slope_slow_, and Slope_very-slow_. Additionally, four global profile features were computed from the full scale range: total area (AUC_total_), maximum and minimum FuzzyEn values (Max_MSE, Min_MSE), complexity index (ComplexityIndex), and profile curvature (Curvature_MSE). All features were evaluated per protocol × *f*_stim_ combination by univariate statistical analysis (Section 2.6).

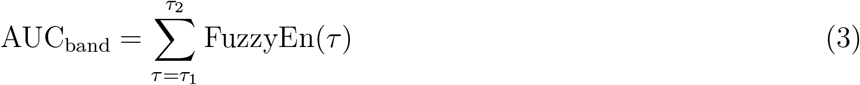

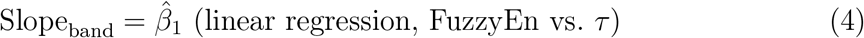

#### 2.5.2. Domain II: FFT Harmonic Analysis

For each recording (*f*_*s*_ = 1953.125 Hz), the DFT was computed over the full length with a rectangular window. For each harmonic *n* = 1, …, 15, the harmonic amplitude ratio is defined as:

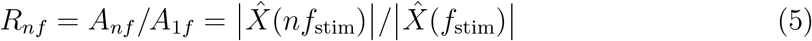

where 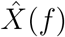 denotes the discrete Fourier transform of the ERG signal evaluated at frequency *f*, and 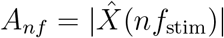 is the spectral amplitude at the *n*-th harmonic. Two additional metrics were computed: the total harmonic distortion (THD), defined as the ratio of the root mean square amplitude of all harmonics *n* ≥ 2 to the fundamental amplitude *A*_1*f*_ [8]:

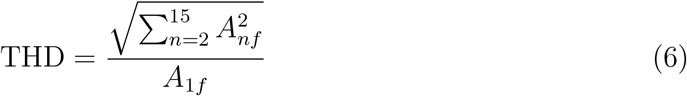

and the narrow-band signal-to-noise ratio (band_snr), defined as the ratio of the spectral power at *f*_stim_ to the mean spectral power in a surrounding noise band excluding the harmonic components [32]. Representative signals and the harmonic spectrum are illustrated in Figure 2, panel A.

#### 2.5.3. Domain III: Time-Frequency Wavelet Coherence (WTC)

Time-frequency wavelet coherence (WTC) is a measure of correlation between two time series in the time-frequency space, a generalization of classical spectral coherence to the wavelet domain [22, 23]. Given two series *x*(*t*) and *y*(*t*), the WTC is defined as:

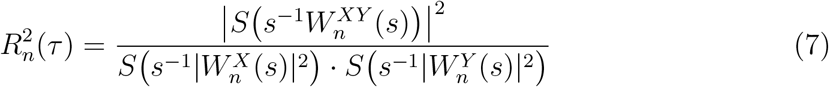

where 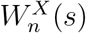 and 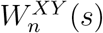 are the wavelet power spectra of *x* and *y* respectively, 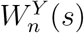 is the cross-wavelet spectrum, *S*(·) is a time-frequency smoothing operator, and *s* is the wavelet scale [22]. 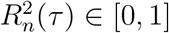, with values close to 1 indicating high coherence between the two signals at scale *s* and time *n*. WTC does not require stationarity and is especially suitable for biological signals with time-varying properties.

In this work, WTC was computed between the preprocessed ERG signal *x*(*t*) and a reference sinusoid *r*(*t*) = sin(2*πf*_stim_*t*) using the Morlet wavelet as basis function (`pycwt.wct`, *δj* = 1*/*12, *ω*_0_ = 6) [22]. The primary extracted metric was the temporal mean of WTC within a ±20% band around *f*_stim_:

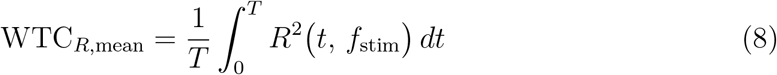

This metric quantifies the global consistency of coupling between the retinal response and the sinusoidal stimulus throughout the recording, independently of the response amplitude [23, 24]. Wavelet coherence has previously been used to quantify stimulus-response coupling in evoked electrophysiological signals [33, 34] and to characterize neural synchronization in pathological states [35]. Figure 2, panel B illustrates the instantaneous WTC for representative subjects from each group. Unlike the other analytical domains, WTC metrics were computed per eye independently and were not bilaterally averaged, given that averaging signals from both eyes prior to coherence computation could cancel inter-ocular phase differences, compromising the estimation of stimulus-response synchrony [22]. Of the two computed metrics—right eye (WTC_*R*,mean_) and left eye (WTC_*L*,mean_)—only the right eye showed significant differences between groups (*p* = 0.0023, |*δ*| = 0.531, large effect), while the left eye did not reach statistical significance (*p* = 0.394, |*δ*| = 0.150, trivial effect). Therefore, WTC_*R*,mean_ was selected as the candidate biomarker for the classification analysis.

#### 2.5.4. Domain IV: Inter-Cycle Sample Entropy

Sample entropy (SampEn) was introduced by Richman and Moorman [36] as a measure of time-series irregularity that is robust to signal length and less sensitive to noise than approximate entropy (ApEn) [37]. For a time series {*x*_*i*_} of length *N*, SampEn is defined as:

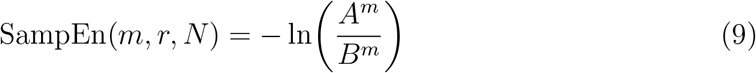

where *B*^*m*^ is the number of template pairs of length *m* that match within a tolerance *r*, and *A*^*m*^ the number of pairs that continue matching when the template is extended to length *m* + 1. A larger SampEn value indicates greater irregularity or complexity of the signal.

Whereas MSFuzzyEn quantifies complexity across multiple temporal scales by progressively coarse-graining the signal, SampEn provides a complementary single-scale measure of irregularity that is sensitive to the local temporal structure of the signal. SampEn has demonstrated its utility for quantifying alterations in signal irregularity in electrophysiological recordings across various neurological conditions [36, 20], including EEG signals in AD [18].

In standard SampEn, template vectors are constructed with a lag of one sample (*τ* = 1), which in a periodic stimulus paradigm captures both the intrinsic variability of the retinal response and the periodicity imposed by the stimulus, conflating two distinct sources of irregularity. To isolate the cycle-to-cycle consistency of the retinal response independently of stimulus periodicity, we introduce a methodological variant in which SampEn is computed using an inter-cycle lag 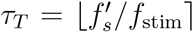, where 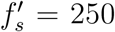 Hz is the decimated sampling rate and *f*_stim_ is the sinusoidal stimulation frequency. This lag corresponds to exactly one full stimulus cycle, so that template vectors compare signal segments separated by one complete retinal response cycle rather than consecutive samples, quantifying irregularity between consecutive cycles independently of the periodicity imposed by the stimulus. The resulting metric, SampEn_*T*_, is defined as:

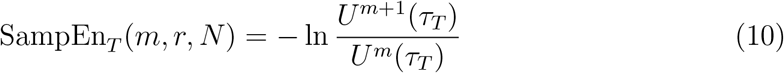

where *U*^*m*^(*τ*_*T*_) is the number of template vector pairs of length *m* within tolerance *r* when vectors are constructed with lag *τ*_*T*_, and *N* is the signal length. Parameters used for SampEn_*T*_ : *m* = 2, *r* = 0.2 *σ*(*x*), where *σ*(*x*) is the standard deviation of the preprocessed decimated ERG signal *x* at the corresponding stimulation frequency *f*_stim_.

#### 2.5.5. Domain V: Oscillatory Potentials (OPs) and DWT

Oscillatory potentials (OPs) are high-frequency oscillations (60–160 Hz) superimposed on the b-wave of the photopic ERG, generated primarily by amacrine cell circuits of the inner retina [9]. OPs are detectable with portable skin-electrode ERG devices including the RETeval system used in this study [38, 39]. Their selective analysis requires time-frequency decomposition methods that preserve the temporal localization of components, which is not possible with classical bandpass filtering [14].

The discrete wavelet transform (DWT) decomposes a signal into time-localized frequency subbands by correlation with a basis function (*mother wavelet*) at multiple resolution scales [40]:

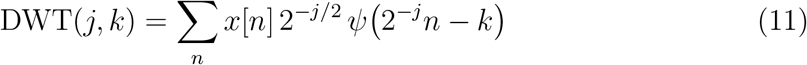

where *j* is the decomposition level, *k* the temporal location, and *ψ* the mother wavelet function. Energy in each subband is calculated as:

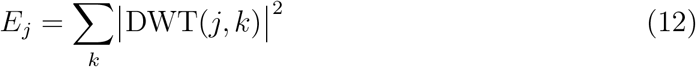

DWT was introduced for ERG OP analysis by Gauvin et al. [13, 14], who demon-strated that energy in the subbands centered at 20, 40, 80, and 160 Hz selectively quantifies contributions from the ON/OFF pathways and from fast and slow OPs, with greater diagnostic sensitivity than classical amplitude analysis. This approach has subsequently been applied to the ERG in neurodevelopmental disorders such as autism and ADHD [15], demonstrating its ability to detect alterations in the inner retina inaccessible to conventional analysis. DWT has also demonstrated superiority over bandpass filtering for OP separation in short-duration electrophysiological signals [16].

In this work, DWT was applied to the averaged signals generated by the RETeval device (ISCEV photopic protocol, *f*_stim_ = 3.433 Hz), using the `db4` wavelet at decomposition level 5, whose detail subband *d*_5_ covers the OP band (≈61–122 Hz). Energy in that subband was calculated as:

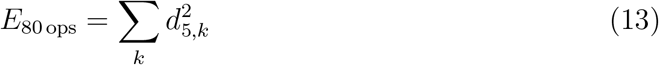

Additionally, the amplitude and latency of peaks OP1, OP2, and OP3 were computed by local maxima detection in the temporal window [15–80] ms, and the amplitude sum OP_amp_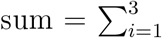 *A*_OP*i*_ as a global measure of inner retinal oscillatory activity. The OP signals are illustrated in Figure 5.

### 2.6. Statistical Analysis

### 2.6.1. Statistical preprocessing

For each metric and protocol × *f*_stim_ combination, values from both eyes were averaged per subject before any statistical comparison, yielding a single representative value per individual (*n*_*C*_ = 26, *n*_*A*_ = 20). This bilateral averaging reduces intra-subject variability and is consistent with standard practice in visual electrophysiology studies [25]. An exception applies to the wavelet time-frequency coherence metric (WTC_*R*,mean_): coherence was computed independently for each eye and treated as two separate candidate metrics in the statistical analysis, rather than averaged bilaterally.

### 2.6.2. Univariate comparisons

Given that normality cannot be assumed in distributions of metrics derived from electrophysiological signals of moderate length [36], the two-sided non-parametric Mann-Whitney *U* test was used to compare each metric between the Controls and AD groups, separately for each protocol × *f*_stim_ combination. The *U* statistic was calculated as:

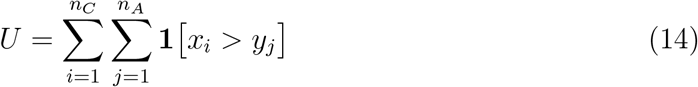

where *x*_*i*_ and *y*_*j*_ are the metric values for subject *i* of the Control group and subject *j* of the AD group, respectively.

#### 2.6.3. Effect size

Cliff’s *δ* [41] was used as the effect size measure, a non-parametric estimator defined as:

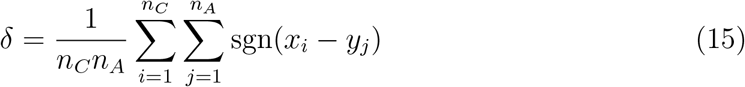

where *δ* ∈ [−1, 1], with positive values indicating predominance of the AD group over Controls. Interpretation thresholds followed Cliff’s criteria [41]: trivial |*δ*| *<* 0.147, small |*δ*| ∈ [0.147, 0.330), medium |*δ*| ∈ [0.330, 0.474), large |*δ*| ≥ 0.474.

#### 2.6.4. Correction for multiple comparisons

To control the false discovery rate (FDR) arising from the multiple comparisons performed, the Benjamini-Hochberg correction [42] was applied per (protocol × metric) family, grouping all stimulation frequencies within each family. For the MSFuzzyEn domain, all 14 derived metrics were included simultaneously in the correction, comprising band-specific AUC and slope metrics as well as global profile features (AUC_total_, Max_MSE, Min_MSE, ComplexityIndex, Curvature_MSE, and wavelet_entropy). The correction was applied per protocol × metric family, grouping the *p*-values from all stimulation frequencies within each protocol × metric combination. The resulting *q*-values were interpreted with a significance threshold of *q <* 0.05. This correction scheme is more conservative than applying FDR globally across all comparisons, preserving FDR control within each family of related hypotheses.

### 2.7. Multivariate Classification

#### 2.7.1. Candidate biomarker selection

The seven candidate biomarkers that reached statistical significance after FDR correction (*q <* 0.05) were considered as classifier candidates, excluding OP1_*time*_ due to its bimodal distribution in the control group (Figure 5B), which compromises the validity of a linear classifier based on this metric.

#### 2.7.2. Search strategy

An exhaustive search was performed over the 2^7^ − 1 = 127 possible combinations of 1 to 7 candidate biomarkers. For each combination, a classifier was trained and evaluated by leave-one-out cross-validation (LOO-CV), which at each fold uses *n* − 1 subjects for training and the remaining subject for evaluation, iterating over all *n* = 46 subjects. LOO-CV is the most appropriate validation scheme for small datasets, as it maximizes available training data at each fold and produces performance estimates with minimum bias for each individual combination [30]. However, selecting the best-performing combination among 127 candidates based on LOO-CV performance introduces an optimistic bias in the reported AUC, since LOO-CV does not account for the multiplicity of the model search. Additionally, candidate biomarkers were pre-selected using univariate statistics of metrics within the same cohort. Therefore, the reported classifier performance should be interpreted as a preliminary upper bound requiring external validation in an independent cohort before any clinical conclusion can be drawn.

#### 2.7.3. Classifier

Logistic regression with *ℓ*_2_ regularization (*C* = 1.0, `scikit-learn` [43]) was used:

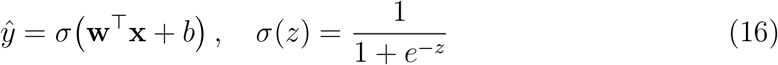

where **x** is the vector of standardized biomarkers, **w** the weight vector, and *b* the bias. Standardization (zero mean, unit standard deviation) was computed exclusively on the training data of each fold and applied to the test observation, preventing data leakage. Logistic regression was selected for its interpretability, its resistance to overfitting with small samples via regularization, and its widespread use in clinical biomarker classification [30].

#### 2.7.4. Optimal classifier selection criterion

In the context of AD screening, the primary objective is the detection of all AD patients, minimizing false negatives—AD individuals misclassified as controls who would not receive timely clinical follow-up. Therefore, sensitivity was prioritized as the primary clinical metric, while specificity was set as a minimum constraint (Spec ≥ 0.80) to avoid an excessive burden of unnecessary diagnostic referrals.

Under this constraint, two classifiers with complementary profiles were identified:

- **Parsimonious classifier (3 candidate biomarkers):** identified from an exhaustive search over all 2^7^ − 1 = 127 possible subsets of the seven candidate biomarkers, by selecting the combination that maximized LOO-CV AUC under the constraint Spec ≥ 0.80 and was consistent with the parsimony rule imposed by the sample size (Section 2.7.5). No domain-independence constraint was imposed during the search. The selected combination spans three independent signal domains (FFT harmonic analysis, wavelet coherence, and MSFuzzyEn), a property of the result rather than a constraint of the search.
- **High-sensitivity classifier (7 candidate biomarkers):** the full set of seven candidate biomarkers selected by maximizing sensitivity at the cost of a moderate reduction in specificity. This classifier is appropriate in population screening scenarios where the clinical cost of a false negative—an undetected AD patient— substantially exceeds that of a false positive.

For both classifiers, the following are reported: sensitivity, specificity, accuracy (Acc), positive predictive value (PPV), and negative predictive value (NPV) calculated from the LOO-CV predictions, together with the corresponding confusion matrices (Figure 6).

It should be noted that both classifiers were selected based on performance evaluated on the same dataset used for candidate biomarker identification, which may introduce optimistic bias. A nested LOO-CV analysis addressing this limitation is reported in the Limitations section.

#### 2.7.5. Parsimony rule

Following the 10-events-per-predictor rule for logistic regression models [44], the maximum number of candidate biomarkers justified by the sample size (*n*_AD_ = 20) is 2 predictors. However, given that LOO-CV conservatively estimates out-of-sample performance and that the selected candidate biomarkers originate from independent signal domains—reducing collinearity as a consequence of the search result rather than by design—it was considered acceptable to extend the parsimonious classifier to 3 candidate biomarkers, with the understanding that results must be replicated in larger cohorts.

### 2.8. Summary of the processing framework

Figure 1 summarizes the complete proposed processing pipeline, from ERG signal acquisition to final classification, integrating both processing branches (sinusoidal and ISCEV) and the five feature extraction domains described in the previous subsections.

## 3. Results

### 3.1. Population-level ERG signals and biomarker profiles

Figure 3 provides a population-level overview of the ERG signals and biomarker profiles across all subjects, illustrating the between-group differences that underlie the seven significant candidate biomarkers identified in this study. Panel A shows the preprocessed ERG signal at *f*_stim_ = 2.996 Hz for all subjects (individual traces, group median and 95% bootstrap CI); Panel B shows the full ISCEV ERG at *f*_stim_ = 3.433 Hz with schematic annotations of the conventional a-wave, b-wave, and PhNR parameters; Panel C shows the MSFuzzyEn profiles at *f*_stim_ = 9.965 Hz with the fast band (*τ* = 1–5) highlighted, illustrating the AUC_fast_ candidate biomarker; Panel D shows the FFT harmonic spectrum at *f*_stim_ = 2.996 Hz with *R*_14*f*_ highlighted; Panel E shows the oscillatory potentials (60–160 Hz) at *f*_stim_ = 3.433 Hz; and Panel F shows the MSFuzzyEn profiles at *f*_stim_ = 39.860 Hz with the very slow band (*τ* = 18–25) highlighted, illustrating the Slope_very-slow_ candidate biomarker.

**Figure 3:**
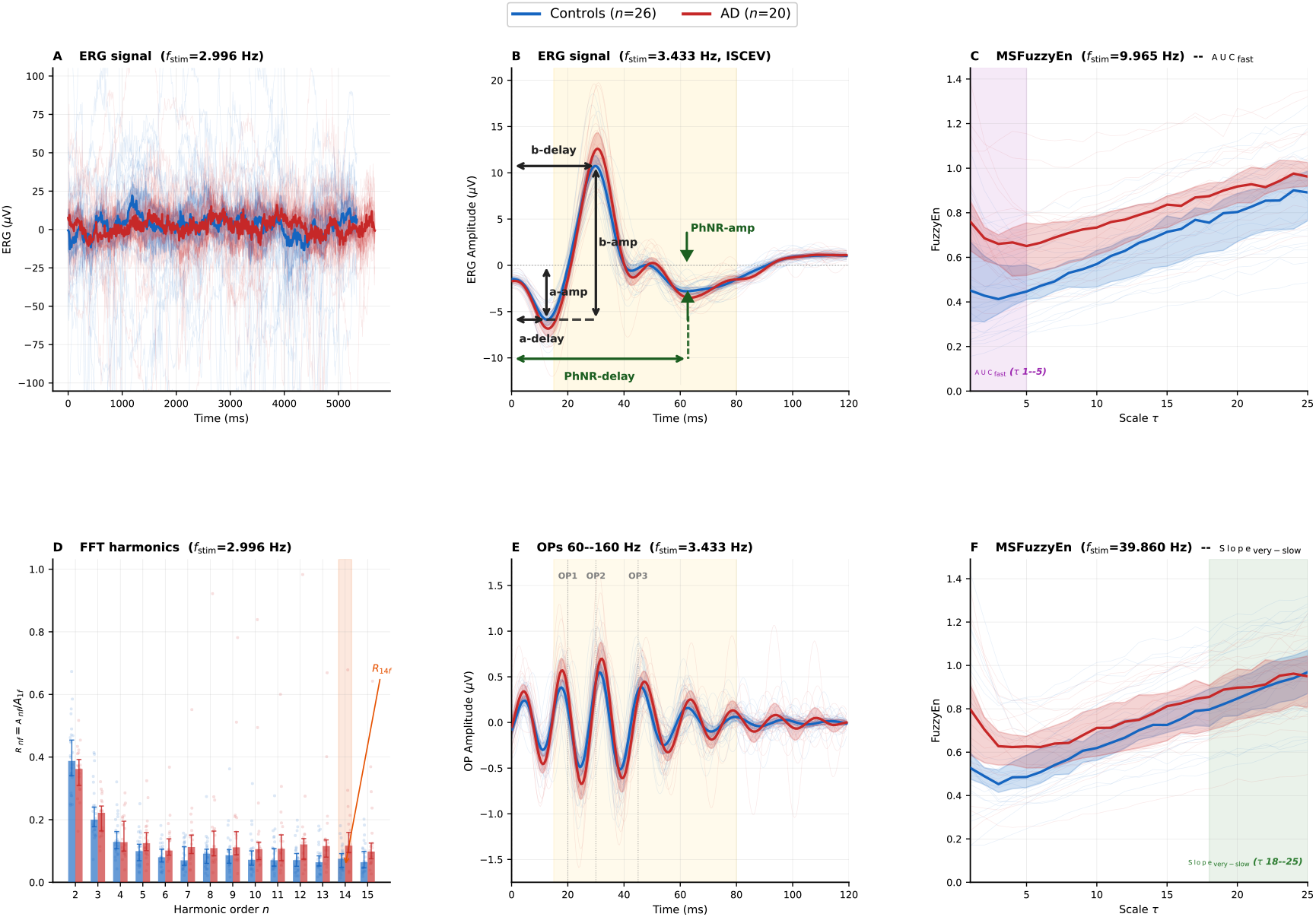
Population-level ERG signals, spectra and complexity profiles — Controls vs AD. **(A)** Preprocessed ERG signal at *f*_stim_ = 2.996 Hz. Individual traces (semitransparent) and group median ± 95% bootstrap CI. **(B)** Full ISCEV ERG at *f*_stim_ = 3.433 Hz (mean ± 95% bootstrap CI). Schematic annotations indicate the measurement of conventional parameters: a-wave amplitude (a-amp) and delay (a-delay), b-wave amplitude (b-amp) and delay (b-delay), and photopic negative response amplitude (PhNR-amp) and delay (PhNR-delay). The yellow-shaded region corresponds to the OP analysis window [15–80] ms. **(C)** MSFuzzyEn profiles at *f*_stim_ = 9.965 Hz. Individual traces and group median ± 95% bootstrap CI. The purple-shaded region (*τ* = 1–5) indicates the fast band over which AUC_fast_ is computed. **(D)** FFT harmonic spectrum at *f*_stim_ = 2.996 Hz. Bars show the group median ± 95% bootstrap CI of *R*_*nf*_ ; individual subject values are shown as semitransparent points. The orange-shaded bar highlights the 14th harmonic (*R*_14*f*_, *f*_14_ ≈ 41.9 Hz). **(E)** Oscillatory potentials (60–160 Hz bandpass filter) at *f*_stim_ = 3.433 Hz (ISCEV protocol). Individual traces and group mean ± 95% bootstrap CI. The yellow-shaded region corresponds to the OP analysis window [15–80] ms. **(F)** MSFuzzyEn profiles at *f*_stim_ = 39.860 Hz. Individual traces and group median ± 95% bootstrap CI. The green-shaded region (*τ* = 18–25) indicates the very slow band over which Slope_very-slow_ is computed. In all panels: blue = Controls (*n* = 26), red = AD (*n* = 20).

### 3.2. Group distribution of significant biomarkers

Figure 4 shows the distribution of the seven biomarkers that reached statistical significance after FDR correction (*q <* 0.05), including the five with large effect size, OP_amp_sum and band_snr with medium effect size. All candidate biomarkers were computed as bilateral averages per subject, except WTC_*R*_ which is associated with the right eye only.

**Figure 4:**
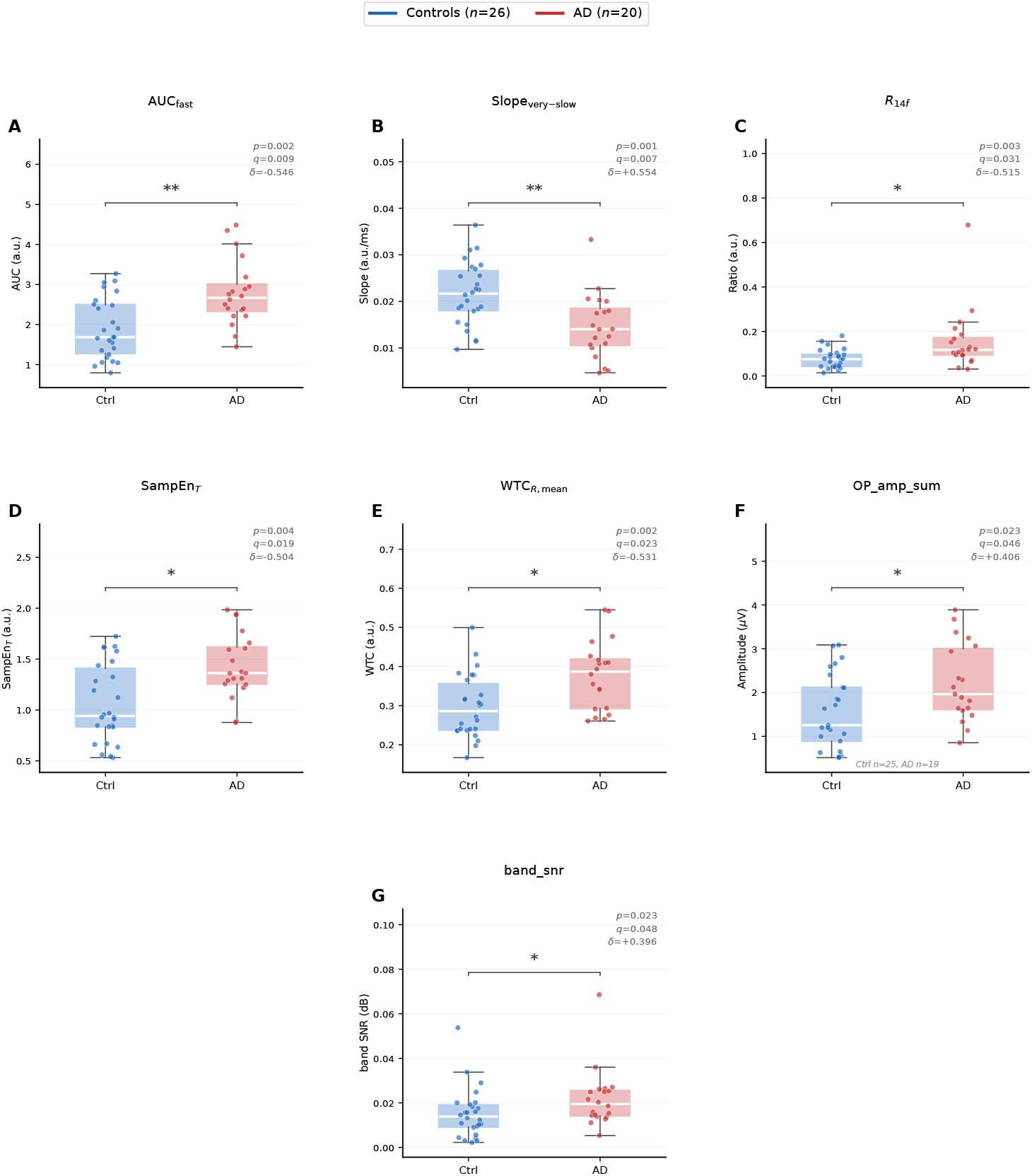
Distribution of the seven significant candidate biomarkers (Mann-Whitney *U*, **FDR-BH** *q <* 0.05) in Controls and AD patients. **(A)** AUC_fast_: area under the MSFuzzyEn profile curve at fast scales (*τ* = 1–5), *f*_stim_ = − 9.965 Hz, 10–50 Hz protocol. AD shows higher values than Controls (*p* = − 0.002; *q* = 0.009; *δ* = −0.546). **(B)** Slope_very-slow_: slope of the MSFuzzyEn profile at very slow scales (*τ* = 18–25), *f*_stim_ = 39.860 Hz. Controls show greater slope than AD (*p* = 0.001; *q* = 0.007; *δ* = +0.554). **(C)** *R*_14*f*_ : spectral amplitude ratio at the 14th harmonic (*f*_14_ ≈ 41.9 Hz), *f*_stim_ = 2.996 Hz, 1–10 Hz protocol. AD shows higher ratio than Controls (*p* = 0.003; *q* = 0.031; *δ* = −0.515). **(D)** Temporal sample entropy (SampEn_*T*_), *f*_stim_ = 9.965 Hz, 10–50 Hz protocol. AD shows higher entropy than Controls (*p* = 0.004; *q* = 0.019; *δ* = −0.504). **(E)** WTC_*R*,mean_: mean time-frequency wavelet coherence, *f*_stim_ = 2.996 Hz, 1–10 Hz protocol. AD shows higher coherence than Controls (*p* = 0.002; *q* = 0.023; *δ* = − 0.531). **(F)** OP_amp_sum: sum of amplitudes of oscillatory potentials OP1, OP2, and OP3, *f*_stim_ = 3.433 Hz, ISCEV photopic protocol. AD shows greater total amplitude than Controls (*p* = 0.023; *q* = 0.046; *δ* = +0.406; *n* = 25 Controls, *n* = 19 AD due to missing data in two subjects). **(G)** band_snr: narrowband signal-to-noise ratio at *f*_stim_ = 9.965 Hz, 10–50 Hz protocol. AD shows higher SNR than Controls (*p* = − 0.023; *q* = 0.048; *δ* = +0.396). In all panels: blue = Controls (*n* = 26), red = AD (*n* = 20) unless otherwise noted. Boxes show median and interquartile range (IQR); whiskers extend to 1.5×IQR. Individual points are shown with horizontal jitter. Significance bars: * *q <* 0.05; ** *q <* 0.01; *** *q <* 0.001. The coefficient *δ* corresponds to Cliff’s delta (Controls − AD). *q*-values corrected by Benjamini-Hochberg FDR over the full set of candidate biomarkers.

#### 3.2.1. AUC_fine_

AUC_fine_ at *f*_stim_ = 9.965 Hz (10–50 Hz protocol) was significantly higher in AD than in Controls (*p* = 0.0017, *q* = 0.009, |*δ*| = 0.546, large effect; median Ctrl: 1.68; median AD: 2.67; Figure 4A). Both group distributions show considerable overlap in their interquartile ranges, although the AD median consistently exceeds that of Controls. The MSFuzzyEn profile of the representative subject from each group at this frequency is illustrated in Figure 2C, where the shaded region (*τ* = 1–5) is the fast scales band over which AUC_fast_ is calculated; the shaded area between both curves reflects the between-group difference (Ctrl: 1.68 a.u.; AD: 2.62 a.u.).

#### 3.2.2. Slope_coarse_

Slope_coarse_ at *f*_stim_ = 39.860 Hz (10–50 Hz protocol) was significantly larger in Controls than in AD (*p* = 0.0015, *q* = 0.007, |*δ*| = 0.554, large effect; median Ctrl: 0.022; median AD: 0.014; Figure 4B). This candidate biomarker had the largest effect size of all identified candidate biomarkers (|*δ*| = 0.554). The AD group distribution shows greater dispersion than Controls, with minimum values near zero. The representative MSFuzzyEn profile for each group is illustrated in Figure 3F, where dashed lines show the linear regression in the very slow scales band (*τ* = 18–25) (Ctrl: 0.0219; AD: 0.0140).

#### 3.2.3. R_14f_

*R*_14*f*_ at *f*_stim_ = 2.996 Hz (1–10 Hz protocol) was significantly higher in AD than in Controls (*p* = 0.0031, *q* = 0.031, |*δ*| = 0.515, large effect; median Ctrl: 0.098; median AD: 0.144; Figure 4C). The 14th harmonic of *f*_stim_ = 2.996 Hz corresponds to *f*_14_ = 14 × 2.996 ≈ 41.9 Hz. The Controls group presents a compact distribution with a narrow interquartile range (values concentrated between ≈0.05 and 0.15), while AD exhibits greater dispersion, with individual values extending up to ≈0.7, including an extreme value near 1.0. The normalized amplitude spectrum of the representative subject from each group is shown in Figure 2A, with the inset magnifying the band around *f*_14_.

#### 3.2.4. SampEn_*T*_

SampEn_*T*_ at *f*_stim_ = 9.965 Hz (10–50 Hz protocol) was significantly higher in AD than in Controls (*p* = 0.0038, *q* = 0.019, |*δ*| = 0.504, large effect; median Ctrl: 0.942; median AD: 1.362; Figure 4D). The Controls group shows greater dispersion than AD, with a wider interquartile range and individual values extending down to ≈0.53. The AD group presents a more compact distribution shifted towards higher values, with all individual values above 0.88.

#### 3.2.5. WTC_*R*,mean_

WTC_*R*,mean_ at *f*_stim_ = 2.996 Hz (1–10 Hz protocol) was significantly higher in AD than in Controls (*p* = 0.0023, *q* = 0.023, |*δ*| = 0.531, large effect; median Ctrl: 0.286; median AD: 0.387; Figure 4E). Unlike the other candidate biomarkers, this metric corresponds exclusively to the right eye, given that the left eye did not reach statistical significance (*p* = 0.394, |*δ*| = 0.150, trivial effect; see Section 2.5.3). The Controls group shows greater dispersion, with individual values ranging from ≈0.20 to ≈0.50, while AD exhibits a more compact distribution shifted towards higher values, with most individual values concentrated between 0.30 and 0.42. The separation between medians is 0.101 units. The instantaneous wavelet coherence WTC_*R*_(*t*) of the representative subject from each group is shown in Figure 2B, lower panel, with dashed lines indicating the temporal mean per group (AD: 0.388; Ctrl: 0.272).

#### 3.2.6. OP*_*amp*_*sum

OP_amp_sum at *f*_stim_ = 3.433 Hz (ISCEV photopic protocol) was significantly higher in AD than in Controls (*p* = 0.0229, *q* = 0.046, |*δ*| = 0.406, medium effect; median Ctrl: 1.256 *µ*V; median AD: 1.965 *µ*V; Figure 4F). Both groups present distributions with considerable overlap; the AD group exhibits a wider interquartile range (IQR: 1.387 *µ*V), with individual values reaching up to 3.888 *µ*V, while Controls present lower central dispersion (IQR: 1.225 *µ*V), with a maximum value of 3.088 *µ*V. *E*_80 ops_ at the same frequency showed a slightly larger effect size (*p* = 0.0153, *q* = 0.031, |*δ*| = 0.423, medium effect) and is reported in Table 3, although it was not included in the figure for consistency with the high-sensitivity classifier.

### 3.3. Conventional ISCEV ERG Parameters

To verify that the candidate biomarkers identified in this study reflect selective alterations in retinal temporal processing rather than global differences in retinal response, conventional ISCEV parameters — a-wave and b-wave amplitude and implicit time, and PhNR amplitude and implicit time — were compared between groups using bilateral averages per subject, Mann-Whitney *U* tests, and Benjamini-Hochberg FDR correction per parameter family across stimulation frequencies, following the same statistical procedure applied to all other metrics in this study. None of the conventional parameters showed significant between-group differences after FDR correction (all *q >* 0.09; Table 2, *n* = 26 Controls, *n* = 20 AD), confirming that the global morphology of the photopic ERG is comparable between groups in this cohort. This finding indicates that the candidate biomarkers identified in this study capture aspects of retinal temporal processing that are inaccessible to standard clinical ERG analysis.

**Table 1:** Participant characteristics. Data as median (IQR) or *n*.

| Characteristic | Controls ($n=26$) | AD ($n=20$) | $p$ |
| --- | --- | --- | --- |
| Age (years) | 66 [60–74] | 70 [65–76] | 0.28 |
| Sex (M/F) | 2/24 | 9/11 | — |

**Table 2:** Conventional ISCEV ERG parameters. Bilateral average per subject, Mann-Whitney *U* test, Benjamini-Hochberg FDR correction per parameter family across stimulation frequencies. Values are medians. None reached statistical significance (all *q >* 0.09). *n* = 26 Controls, *n* = 20 AD, except PhNR at *f* = 1.963 Hz (*n* = 25 Controls).

| Parameter | $f_{\text{stim}}$ (Hz) | Controls | AD | $p$ | $q$ | $\delta$ |
| --- | --- | --- | --- | --- | --- | --- |
| a-wave amplitude ($\mu\text{V}$) | 1.963 | 4.68 | 5.94 | 0.053 | 0.094 | -0.338 |
| a-wave amplitude ($\mu\text{V}$) | 3.433 | 5.86 | 6.89 | 0.094 | 0.094 | -0.292 |
| a-wave implicit time (ms) | 1.963 | 12.05 | 11.64 | 0.313 | 0.419 | +0.177 |
| a-wave implicit time (ms) | 3.433 | 12.74 | 12.89 | 0.419 | 0.419 | -0.142 |
| b-wave amplitude ($\mu\text{V}$) | 1.963 | 21.61 | 21.87 | 0.485 | 0.485 | -0.123 |
| b-wave amplitude ($\mu\text{V}$) | 3.433 | 16.87 | 19.43 | 0.086 | 0.172 | -0.300 |
| b-wave implicit time (ms) | 1.963 | 30.26 | 30.15 | 0.542 | 0.542 | +0.108 |
| b-wave implicit time (ms) | 3.433 | 30.07 | 30.73 | 0.099 | 0.198 | -0.288 |
| PhNR amplitude ($\mu\text{V}$) | 1.963 | 3.00 | 5.31 | 0.077 | 0.153 | -0.312 |
| PhNR amplitude ($\mu\text{V}$) | 3.433 | 2.58 | 2.22 | 0.731 | 0.731 | +0.062 |
| PhNR implicit time (ms) | 1.963 | 50.18 | 49.15 | 0.900 | 0.900 | -0.024 |
| PhNR implicit time (ms) | 3.433 | 50.28 | 45.36 | 0.283 | 0.565 | +0.188 |

### 3.4. Oscillatory Potentials

Oscillatory potentials (OPs) were extracted and quantified by discrete wavelet transform (DWT, `db4` wavelet, decomposition level 5) applied to the ISCEV photopic ERG averaged signal at *f*_stim_ = 3.433 Hz, whose detail subband *d*_5_ covers the OP band (≈61–122 Hz). For visualization purposes, the signals shown in Figure 5 were obtained with a zero-phase Butterworth bandpass filter [60–160] Hz applied to the same averaged signal. Figure 5 summarizes the main OP analysis findings.

**Figure 5:**
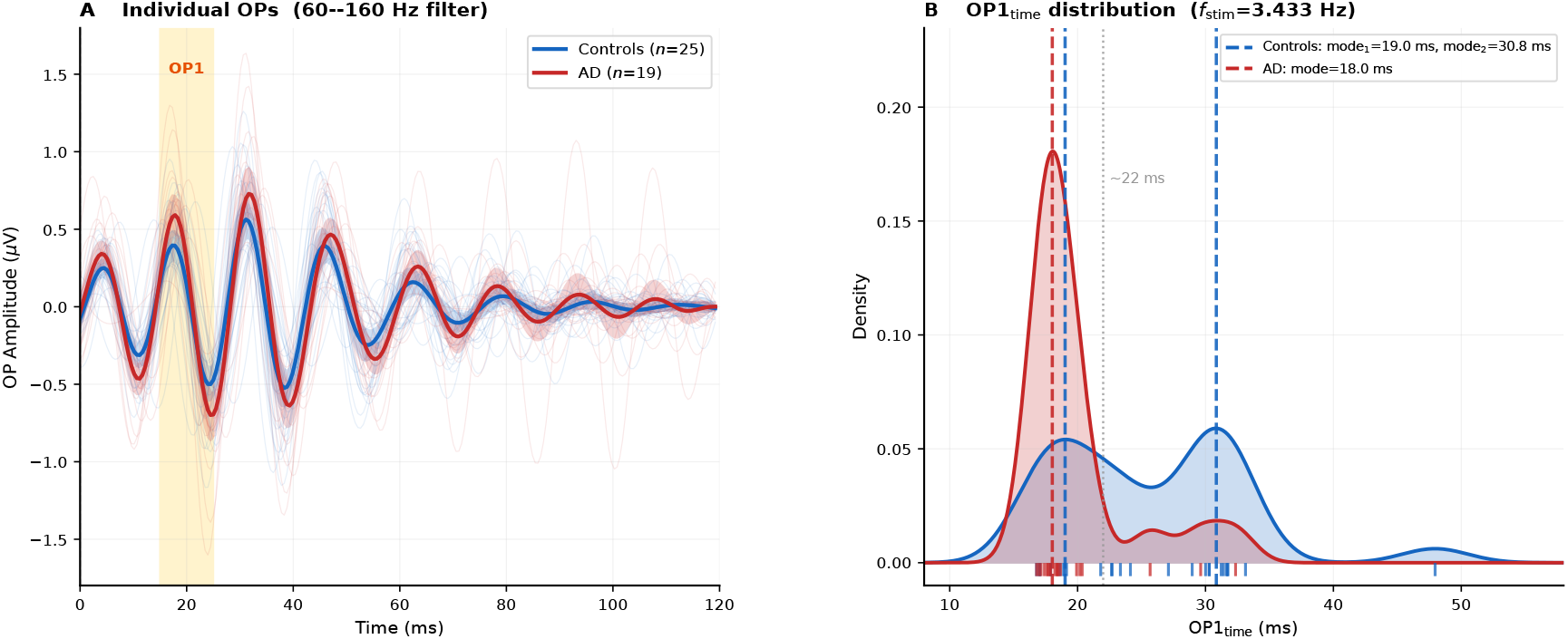
Oscillatory potentials (OPs) of the ISCEV photopic ERG at *f*_stim_ = 3.433 Hz. **(A)** Individual OP signals (60–160 Hz bandpass filter) for each subject, superimposed by group: Controls (blue, *n* = 25) and AD (red, *n* = 19). Thick line indicates the group mean ± 95% bootstrap CI. The yellow-shaded region highlights the OP1 window (15–25 ms). **(B)** Estimated density distribution (KDE, bandwidth= 0.35) of OP1 implicit time (OP1_*time*_). Controls exhibit a bimodal distribution with two modes at ∼19.0 ms and ∼30.8 ms (vertical dashed blue lines); AD presents a concentrated unimodal distribution with mode at ∼18.4 ms (vertical dashed red line). The grey dotted line indicates the approximate cut-off between modes (∼22 ms). Tick marks at the base represent individual values (rug plot). In all panels: *n* = 25 Controls and *n* = 19 AD.

Figure 5A shows individual OP signals from all subjects superimposed by group. The yellow-shaded region highlights the OP1 window [15–25] ms. Individual traces of the AD group (red) present visibly larger amplitudes than those of the Controls group (blue) within that window, with greater inter-subject variability. The group mean (thick line) confirms this difference. Consistent with the formal statistical analysis presented in Table 2, neither the a-wave nor the b-wave showed significant between-group differences in amplitude or implicit time (all *q >* 0.09), confirming that the global morphology of the photopic ERG is comparable between AD and Controls in this cohort. This finding contrasts with the differences observed in the OPs, which reflect selective activity of inner retinal circuits (Figure 3B,E).

Figure 5B shows the estimated density distribution (KDE) of OP1_*time*_ for both groups. The Controls group presents a bimodal distribution with two clearly differentiated modes around ∼19.0 ms and ∼30.8 ms, while the AD group presents a unimodal distribution concentrated around ∼18.4 ms. The grey dotted line indicates the approximate cut-off between modes (∼22 ms). The difference in OP1_*time*_ between groups was statistically significant and had the largest effect size among all OP biomarkers (*p* = 0.0008, *q* = 0.008, |*δ*| = 0.512, large effect). Nevertheless, this metric was not included in the classifier or among the candidate biomarkers, given that the bimodal distribution observed in the control group (Figure 5B) violates the assumptions of a linear classifier and complicates the interpretation of any decision threshold based on this variable.

### 3.5. Summary of univariate results

Table 3 summarizes the seven candidate biomarkers that reached statistical significance after FDR correction (*q <* 0.05), ordered by effect size. The significant metrics span five independent signal domains (MSFuzzyEn, WTC, FFT, SampEn, and OPs-ISCEV) and three stimulation frequencies (2.996, 9.965, and 39.860 Hz). Five candidate biomarkers showed large effect (|*δ*| ≥ 0.474) and two showed medium effect. AD patients showed higher values than Controls for AUC_fast_, WTC_*R*,mean_, *R*_14*f*_, SampEn_*T*_, OP_amp_sum, and band_snr, while Controls showed a steeper Slope_very-slow_ than AD patients. The three candidate biomarkers of the parsimonious classifier are indicated in bold.

**Table 3:**
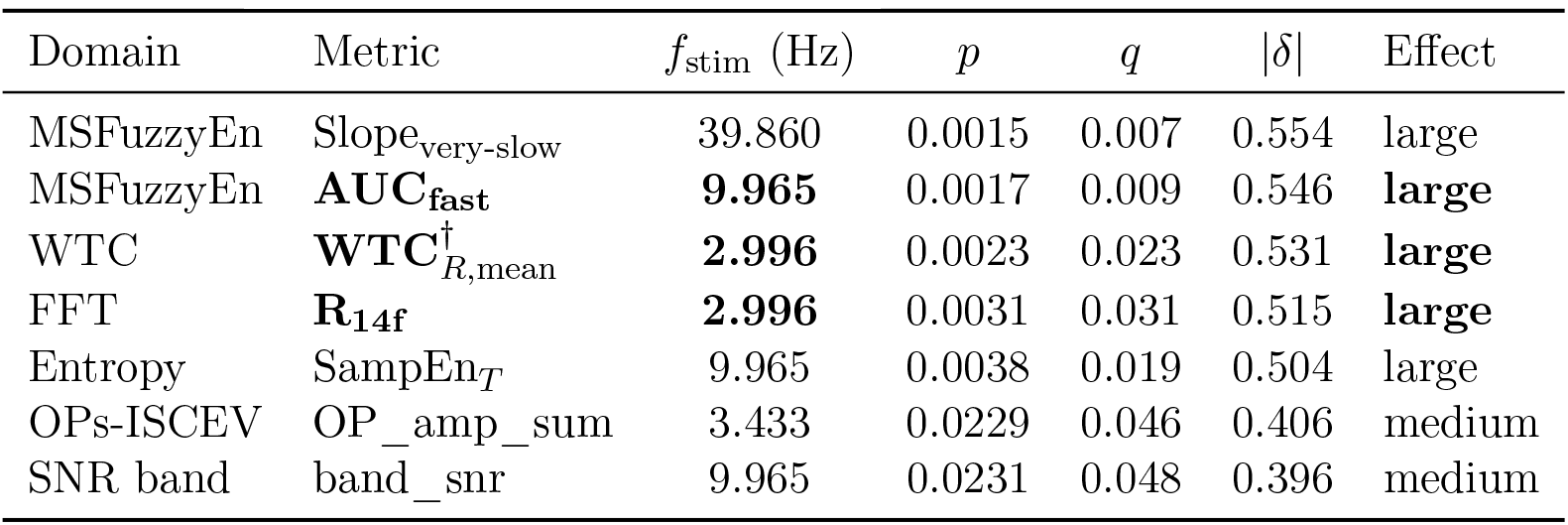
Univariate results. Bilateral average per subject, Mann-Whitney *U* test, FDR correction per protocol × metric family. *δ*: Cliff’s delta (positive: AD > Ctrl). **Bold**: candidate biomarkers of the optimal classifier (3 biom.). ^*†*^: right eye exclusively.

### 3.6. Multivariate classification

Given the small sample size (*n* = 46) and the absence of an independent holdout set, the following results should be interpreted as preliminary and hypothesis-generating. Candidate biomarker selection was based on univariate significance in the same dataset used for classifier construction, a limitation that LOO-CV does not fully correct. Re-ported performance estimates should therefore be regarded as optimistic upper bounds pending replication in an independent cohort.

The exhaustive search over 127 combinations identified the combination of three candidate biomarkers from independent domains as the optimal classifier (Spec ≥ 0.80, maximum AUC): AUC_fast_ + *R*_14*f*_ + WTC_*R*,mean_.

**Optimal classifier (3 candidate biomarkers)**, *n* = 46:

- AUC = 0.858
- Sensitivity = 0.700 (14/20 AD correctly classified)
- Specificity = 0.885 (23/26 Ctrl correctly classified)
- Acc = 0.804; PPV = 0.824; NPV = 0.793

**High-sensitivity classifier (7 candidate biomarkers)**, *n* = 44: adds Slope_very-slow_, SampEn_*T*_, OP_amp_sum, and band_snr:

- AUC = 0.825
- Sensitivity = 0.789 (15/19 AD correctly classified)
- Specificity = 0.840 (21/25 Ctrl correctly classified)
- Acc = 0.818; PPV = 0.789; NPV = 0.840
- Note: the dataset reduces to *n* = 44 due to missing data in one or more additional candidate biomarkers (Ctrl= 25, AD= 19)

The ROC curves of both classifiers, their confusion matrices, and the comparison with the individual ROC curves of each candidate biomarker in the optimal classifier are presented in Figure 6. Panel A shows that the 3-candidate biomarker optimal classifier (AUC= 0.858, Spec= 0.885, solid green line, circular marker) outperforms the 7-candidate biomarker high-sensitivity classifier (AUC= 0.825, Spec= 0.840, dashed red line, diamond marker) in terms of AUC and specificity, while the 7-candidate biomarker classifier achieves higher sensitivity (0.789 vs 0.700). Panel B shows that the 3-candidate biomarker multivariate model (AUC= 0.858) outperforms each individual candidate biomarker evaluated in isolation: AUC_fast_ (AUC= 0.773), *R*_14*f*_ (AUC= 0.758), and WTC_*R*,mean_ (AUC= 0.765), confirming the complementarity of the three analytical domains.

**Figure 6:**
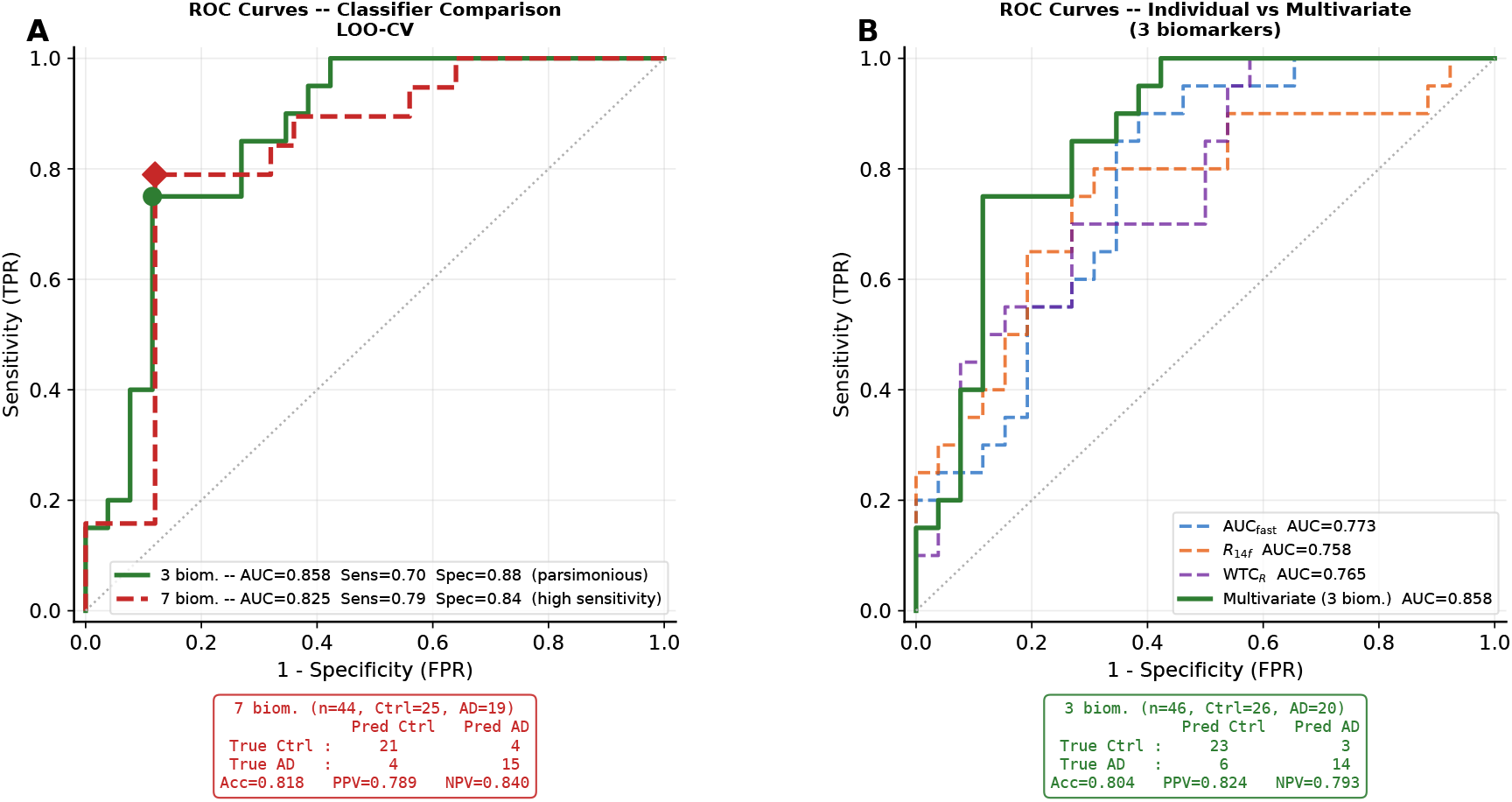
ROC curves and confusion matrices of the ERG-Alzheimer classifiers evaluated by LOO-CV. **(A)** Comparison between the parsimonious classifier (3 candidate biomarkers: AUC_fast_, *R*_14*f*_, WTC_*R*,mean_; solid green line) and the high-sensitivity classifier (7 candidate biomarkers: AUC_fast_, Slope_very-slow_, *R*_14*f*_, SampEn_*T*_, WTC_*R*,mean_, OP_amp_sum, band_snr; dashed red line). The parsimonious classifier achieves AUC= 0.858, Sensitivity= 0.700, Specificity= 0.885 (*n* = 46; Ctrl= 26, AD= 20). The high-sensitivity classifier achieves AUC= 0.825, Sensitivity= 0.789, Specificity= 0.840 (*n* = 44; Ctrl= 25, AD= 19). The filled circle (green) indicates the optimal operating point of the parsimonious classifier and the diamond marker (red) that of the high-sensitivity classifier (maximum of Sensitivity − FPR). The lower confusion matrix corresponds to the high-sensitivity classifier (Acc= 0.818; PPV= 0.789; NPV= 0.840). **(B)** Comparison between the individual ROC curves of each candidate biomarker in the parsimonious classifier (AUC_fast_: AUC= 0.773; *R*_14*f*_ : AUC= 0.758; WTC_*R*,mean_: AUC= 0.765; dashed lines) and the ROC curve of the combined multivariate model (AUC= 0.858; solid green line). The lower confusion matrix corresponds to the parsimonious classifier (Acc= 0.804; PPV= 0.824; NPV= 0.793). All models were evaluated by leave-one-out cross-validation (LOO-CV). Classifier implemented with *ℓ*_2_-regularised logistic regression (*C* = 1.0, random state= 42).

## 4. Discussion

### 4.1. MSFuzzyEn and multiscale ERG complexity in AD

Amplitude and implicit time analysis of ERG waves constitutes the current clinical standard, but captures only average properties of the retinal response, ignoring the complex temporal structure of the signal cycle by cycle. Multiscale fuzzy entropy (MSFuzzyEn) overcomes this limitation by quantifying signal irregularity at multiple temporal scales simultaneously, being sensitive to both amplitude and phase dynamics of oscillations, unlike conventional power spectral analysis [18].

The validity of this approach for the ERG in AD is directly supported by Araya-Arriagada et al. [17], who demonstrated in the 5xFAD mouse model that conventional amplitude metrics of the microERG showed no between-group differences while multiscale entropy analysis did detect significant differences—precisely the pattern observed in our data, where a-waves and b-waves are comparable between groups (Figure 3B) but entropy metrics are significantly different (Figure 4A,D).

AUC_fast_ quantifies complexity in the fast scale band (*τ* = 1–5, equivalent to 25– 125 Hz), where the between-group separation is maximal (Figure 2C). Slope_very-slow_ captures the hierarchical organization of complexity across scales (*τ* = 18–25, equivalent to 5–7 Hz): a positive slope indicates that the signal increases its complexity at slower scales, a pattern characteristic of healthy physiological signals [31], which flattens significantly in AD (Figure 3F). Both metrics are complementary: AUC_fast_ captures the absolute complexity level at high frequency, while Slope_very-slow_ captures the multiscale organization of the signal. The systematic application of MSFuzzyEn to the human ERG in a clinical context constitutes an original contribution of the present work.

### 4.2. FFT harmonic analysis and retinal nonlinearity

Amplitude analysis of the sinusoidal ERG in the frequency domain is conventionally limited to the fundamental frequency (*f*_stim_) and the second harmonic (2*f*_stim_), ignoring the information contained in higher harmonics [8, 9]. The ratio *R*_*nf*_ = *A*_*nf*_ */A*_1*f*_ proposed in this work extends this analysis to harmonics of arbitrary order, allowing quantification of the relative energy of high-frequency nonlinear components that are invisible to conventional amplitude analysis.

*R*_14*f*_ at *f*_stim_ = 2.996 Hz corresponds to the 14th harmonic at ≈ 42 Hz. Its statistical significance (*q* = 0.031, |*δ*| = 0.515, large effect) demonstrates that the AD retina generates proportionally more energy at this high-frequency harmonic than controls, a pattern inaccessible to conventional amplitude and implicit time analysis. This finding is consistent with previous studies documenting greater harmonic distortion in sinusoidal ERGs under conditions of retinal dysfunction [10], where alterations in higher harmonics precede changes in the fundamental.

The complementarity of *R*_14*f*_ with entropy candidate biomarkers is evidenced in the optimal classifier: the three candidate biomarkers originate from independent domains— temporal complexity (MSFuzzyEn), harmonic spectrum (FFT), and time-frequency coherence (WTC)—capturing distinct aspects of the retinal response that cannot be reduced to a single analysis.

### 4.3. Inter-cycle sample entropy and wavelet coherence: irregularity and stimulus-response synchrony

SampEn_*T*_ and WTC_*R*,mean_ quantify complementary properties of the sinusoidal ERG response that conventional analysis does not capture.

Inter-cycle sample entropy with lag 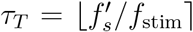 constitutes a specific methodological contribution of this work: rather than quantifying irregularity in absolute time, it directly quantifies the cycle-to-cycle consistency of the ERG response. This approach is more appropriate for signals responding to periodic stimuli than standard sample entropy, as the inter-cycle lag removes the stimulus periodicity from the complexity estimate, capturing exclusively the variability of the response between consecutive cycles [36]. SampEn_*T*_ greater in AD (*q* = 0.019, |*δ*| = 0.504) indicates that the ERG response is less stereotyped cycle by cycle in AD patients than in controls, a pattern consistent with the greater irregularity detected by MSFuzzyEn.

Time-frequency wavelet coherence (WTC) offers a methodologically distinct perspective: rather than quantifying the intrinsic complexity of the signal, it measures the synchrony between the ERG response and the reference sinusoidal stimulus over time. The fundamental advantage of WTC over Fourier spectral coherence is that it does not assume stationarity of the signal—a requirement that the sinusoidal ERG does not strictly meet [12]— and allows detection of changes in stimulus-response coherence with temporal resolution, capturing features inaccessible through Fourier coherence [33, 22]. WTC_*R*,mean_ greater in AD (*q* = 0.023, |*δ*| = 0.531) indicates greater stimulus-response coherence in the right eye of AD patients at *f*_stim_ = 2.996 Hz. The ocular asymmetry— with the left eye not significant (*p* = 0.394, |*δ*| = 0.150)—is a finding that warrants specific investigation in future studies.

### 4.4. Oscillatory potentials and DWT: selective energy extraction

Conventional analysis of oscillatory potentials (OPs) through 60–160 Hz bandpass filtering presents well-documented methodological limitations: it introduces phase distortion, ringing artefacts, and amplitude attenuation that can generate artificial OPs or suppress real OPs [14]. The discrete wavelet transform (DWT) overcomes these limitations by providing time-frequency decomposition with preserved temporal localization, without the artefacts inherent to bandpass filtering [13, 14].

In this work DWT with `db4` wavelet was applied at decomposition level 5, whose subband *d*_5_ covers ≈61–122 Hz, selectively capturing the energy of OPs without contamination from slow waves (a and b). *E*_80 ops_ greater in AD (*q* = 0.031, |*δ*| = 0.423) and OP_amp_sum greater in AD (*q* = 0.046, |*δ*| = 0.406) demonstrate that the oscillatory energy of the inner retina is greater in AD than in controls, while the global ERG morphology (a and b waves) is comparable between groups (Figure 3B). This pattern—selective OP dysfunction with preserved conventional ERG—has been previously documented in other conditions of selective inner retinal dysfunction, such as diabetic retinopathy [14] and neurodevelopmental disorders [15]. Its presence in AD suggests that DWT OP analysis captures inner retinal alterations inaccessible to conventional a and b wave amplitude analysis.

The finding of a bimodal distribution of OP1_*time*_ in Controls (Figure 5B) is methodologically relevant in two respects. First, it illustrates that statistically significant metrics can be inappropriate for linear classification when they violate classifier assumptions, underscoring the importance of evaluating the distribution of candidate biomarkers before incorporating them into classification models. Second, the contrast between the bimodal distribution of Controls—with two modes around ∼19 ms and ∼30.8 ms—and the concentrated unimodal distribution of AD at ∼18.4 ms suggests that OP1 latency may reflect subpopulations of retinal response with distinct temporal dynamics in healthy subjects, a phenomenon that becomes homogenized in AD. This observation, although not explained by the available demographic variables (age and sex), warrants specific investigation in future studies with larger sample sizes and more detailed phenotypic characterization.

### 4.5. Domain complementarity and multivariate classifier

The diagnostic gain of the 3-candidate biomarker optimal classifier (AUC= 0.858) over each individual candidate biomarker (AUC: 0.758–0.773) demonstrates that the three analytical domains capture complementary, non-redundant information about the ERG response in AD (Figure 6B). This complementarity is expected given that the three candidate biomarkers quantify fundamentally distinct signal properties: AUC_fast_ the temporal complexity at high frequency, *R*_14*f*_ the spectral nonlinearity, and WTC_*R*,mean_ the stimulus-response synchrony in the time-frequency domain.

Compared with the PhNR-based classifier reported by Asanad et al. [7] (AUC= 0.84, Sens= 0.87, Spec= 0.82), the optimal classifier of this work achieves comparable performance (AUC= 0.858) with the methodological advantage of using only sinusoidal ERGs, recorded with a portable device without pupil dilation, versus the ISCEV protocol with pharmacological dilation and laboratory equipment required by Asanad et al.

The 7-candidate biomarker high-sensitivity classifier (AUC= 0.825, Sens= 0.789) incorporates medium-effect candidate biomarkers that provide additional information about the retinal response in AD, at the cost of greater model complexity and reduced effective sample size (*n* = 44 vs *n* = 46) due to missing data in some candidate biomarkers. The sensitivity-specificity trade-off between both classifiers (Figure 6A) is relevant in the screening context, where clinical priority determines which profile is more appropriate.

### 4.6. Limitations

The sample size is modest (*n* = 46), which limits statistical power and the stability of LOO-CV estimators. Candidate biomarker selection and classifier evaluation were performed on the same dataset without an independent holdout set, so the reported performance figures should be considered preliminary estimates subject to external validation [30]. To assess the optimism bias introduced by selecting the best-performing combination among 127 candidate models, a nested LOO-CV procedure was additionally performed, in which the inner exhaustive search was conducted independently within each outer fold [30]. This yielded AUC= 0.821, sensitivity= 0.600 and specificity= 0.923 (*n* = 46), representing a modest correction of ΔAUC= 0.037 relative to the standard LOO-CV estimate. The most frequently selected combination in the inner search (42/46 folds) was *E*_80 ops_ + *R*_14*f*_ + Slope_very-slow_, which differs from the optimal classifier identified by the standard procedure, underscoring the instability of model selection in small samples. These findings confirm that the reported classifier performance should be regarded as a preliminary upper bound requiring replication in an independent cohort before clinical translation.

The sex distribution is unbalanced, and no age adjustment was performed in the univariate analysis. Finally, the ocular asymmetry observed in WTC (WTC_*R*,mean_ significant, WTC_*L*,mean_ not significant) has no clear methodological explanation and requires investigation in larger cohorts.

### 4.7. Implications and future work

The proposed multi-domain framework demonstrates that ERG signal analysis extends beyond conventional amplitude and implicit time parameters, extracting diagnostically relevant information through five complementary techniques applied to signals recorded with a non-invasive portable device. Results support the feasibility of using the RETeval device in community-based AD screening protocols, without requiring pupil dilation or specialized laboratory equipment.

Future work should include: (i) replication in larger cohorts with confirmation by molecular biomarkers; (ii) evaluation at preclinical stages (mild cognitive impairment) where screening has the greatest clinical impact; (iii) longitudinal follow-up of the proposed metrics as progression indicators; and (iv) investigation of the ocular asymmetry in WTC and its possible independent diagnostic value.

## 5. Conclusions

We have presented a multi-domain signal analysis framework for the detection of Alzheimer’s disease from full-field sinusoidal ERGs recorded with a non-invasive portable device. The framework integrates five electrophysiological signal processing techniques— multiscale fuzzy entropy (MSFuzzyEn), FFT harmonic analysis, time-frequency wavelet coherence (WTC), inter-cycle sample entropy (SampEn_*T*_), and discrete wavelet transform (DWT) for oscillatory potential extraction—each capturing distinct and independent properties of the ERG signal (Figure 1).

Seven candidate biomarkers showed significant differences between AD and controls after FDR correction, with five achieving large effect size (Table 3, Figure 4): AUC_fast_ (|*δ*| = 0.546, *q* = 0.009), Slope_very-slow_ (|*δ*| = 0.554, *q* = 0.007), *R*_14*f*_ (|*δ*| = 0.515, *q* = 0.031), SampEn_*T*_ (|*δ*| = 0.504, *q* = 0.019), and WTC_*R*,mean_ (|*δ*| = 0.531, *q* = 0.023); and two with medium effect size: OP_amp_sum (|*δ*| = 0.406, *q* = 0.046) and band_snr (|*δ*| = 0.396, *q* = 0.048).

A logistic regression classifier with three candidate biomarkers (AUC_fast_ + *R*_14*f*_ + WTC_*R*,mean_) achieved AUC = 0.858, sensitivity = 70.0%, and specificity = 88.5% in LOO-CV (*n* = 46, Figure 6), a result superior to that of any individual candidate biomarker and consistent with the parsimony constraint imposed by the sample size.

These findings demonstrate that multi-domain signal processing of the sinusoidal ERG reveals signatures of retinal temporal processing dysfunction in AD—increased complexity at fast scales, altered nonlinearity in the gamma band, paradoxical hypersynchronization, and selective hyperexcitability of inner retinal circuits—that are inaccessible to standard clinical analysis and contain substantial diagnostic information for non-invasive AD screening.

## Acknowledgements

This work was funded by ANID (Agencia Nacional de Investigación y Desarrollo) via project Exploración 13220082 granted to LEM and AGP. ERG recordings were carried out at the Ophthalmology Unit of Hospital del Salvador, Santiago, Chile. The authors thank all participating patients and volunteers, as well as the ophthalmological technical staff of Hospital del Salvador.

## Ethical approval

The study was approved by the Institutional Ethics Committee of Universidad de Santiago de Chile (Ethics Report No. 323/2023). All participants signed informed consent.

## Data and code availability

Anonymized metric data and the analysis code can be obtained by contacting the corresponding author.

